# Condition-dependence resolves the paradox of missing plasticity costs

**DOI:** 10.1101/2022.09.30.510277

**Authors:** Stephen P. De Lisle, Locke Rowe

## Abstract

Phenotypic plasticity plays a key role in adaptation to changing environments. However, plasticity is neither perfect nor ubiquitous, implying that fitness costs must limit the evolution of phenotypic plasticity in nature. The measurement of such costs of plasticity has proved elusive; decades of experiments show that fitness costs of plasticity are often weak or nonexistent. Here, we show that this paradox can be at least partially explained by condition-dependence. We develop two models differing in their assumptions about how condition-dependence arises; both models show that variation in condition can readily mask costs of plasticity even when such costs are substantial. This can be shown simply in a model where costly plasticity itself evolves condition-dependence. Yet similar effects emerge from an alternative model where trait expression is condition-dependent. In this more complex model, average condition in each environment and genetic covariance in condition across environments both determine when costs of plasticity can be revealed. Analogous to the paradox of missing trade-offs between life history traits, our models show that variation in condition masks costs of plasticity even when costs exist, and suggests this conclusion may be robust to the details of how condition affects trait expression. Our models demonstrate that condition dependence can also account for the often-observed pattern of elevated plasticity costs inferred in stressful environments, the maintenance of genetic variance in plasticity, and provides insight into experimental and biological scenarios ideal for revealing a cost of phenotypic plasticity.

## Introduction

Phenotypic plasticity occurs when the same genotype produces different phenotypes in response to different local or developmental environments. Plasticity, when adaptive, allows organisms to track an environment-dependent optimum within a single generation, permitting expression of adaptive phenotypes in a new environment and preventing maladaptation in temporally- or spatially-variable environments (Charmantier *et al*. 2008; Chevin *et al*. 2010; Chevin & Lande 2011). For these reasons, plasticity is an important source of phenotypic variation, and the evolution of phenotypic plasticity plays a key role in adaptation to variable or novel environments (Charmantier *et al*. 2008; Phillimore *et al*. 2010; Charmantier & Gienapp 2014; Crozier & Hutchings 2014; Snell-Rood *et al*. 2018; Stamp & Hadfield 2020).

However, no organism or trait is completely plastic, despite the clear benefits of altering development in response to the environment. This implies that fitness costs of plasticity or genetic constraints (“limits”) must play a role in restricting the evolution of plasticity in nature (Callahan *et al*. 2008; Auld *et al*. 2010; Murren *et al*. 2015). Although genetic constraints must play a role in the evolution of all traits, including plasticity, genetic constraints are an insufficient explanation for limited plasticity. Empirically, this is because standing genetic variance is often observed for reaction norms (Scheiner 1993), and theoretically, because unless genetic constraints are complete they simply slow the rate of plasticity evolution (Via & Lande 1985; Van Tienderen 1991). This implies some fitness cost of plasticity must exist (DeWitt *et al*. 1998) - there must be a fitness penalty for the ability to alter development or behavior in response to the environment, that balances with the benefits of tracking an environmentally-dependent trait optimum.

A fitness cost of plasticity is expected to manifest as a reduction in fitness of a plastic genotype compared to a non-plastic genotype that otherwise expresses the same trait value in a given environment (Figure 1A). Empirically, such costs are often investigated in a Genotype x Environment (GxE) experiment in which the same set of genotypes are reared in two environments, followed by a phenotypic selection analysis (Lande & Arnold 1983) in which genotype fitness in a focal environment is regressed against both the trait value expressed in the focal environment as well as well as some measure of plasticity, which depends on trait expression in the second environment (Van Tienderen 1991; DeWitt 1998; DeWitt *et al*. 1998; Scheiner & Berrigan 1998; Van Buskirk & Steiner 2009; Auld *et al*. 2010). A cost of plasticity is expected to manifest as a negative partial regression coefficient for the measure of a genotype’s plasticity and fitness.

**Figure 1.**
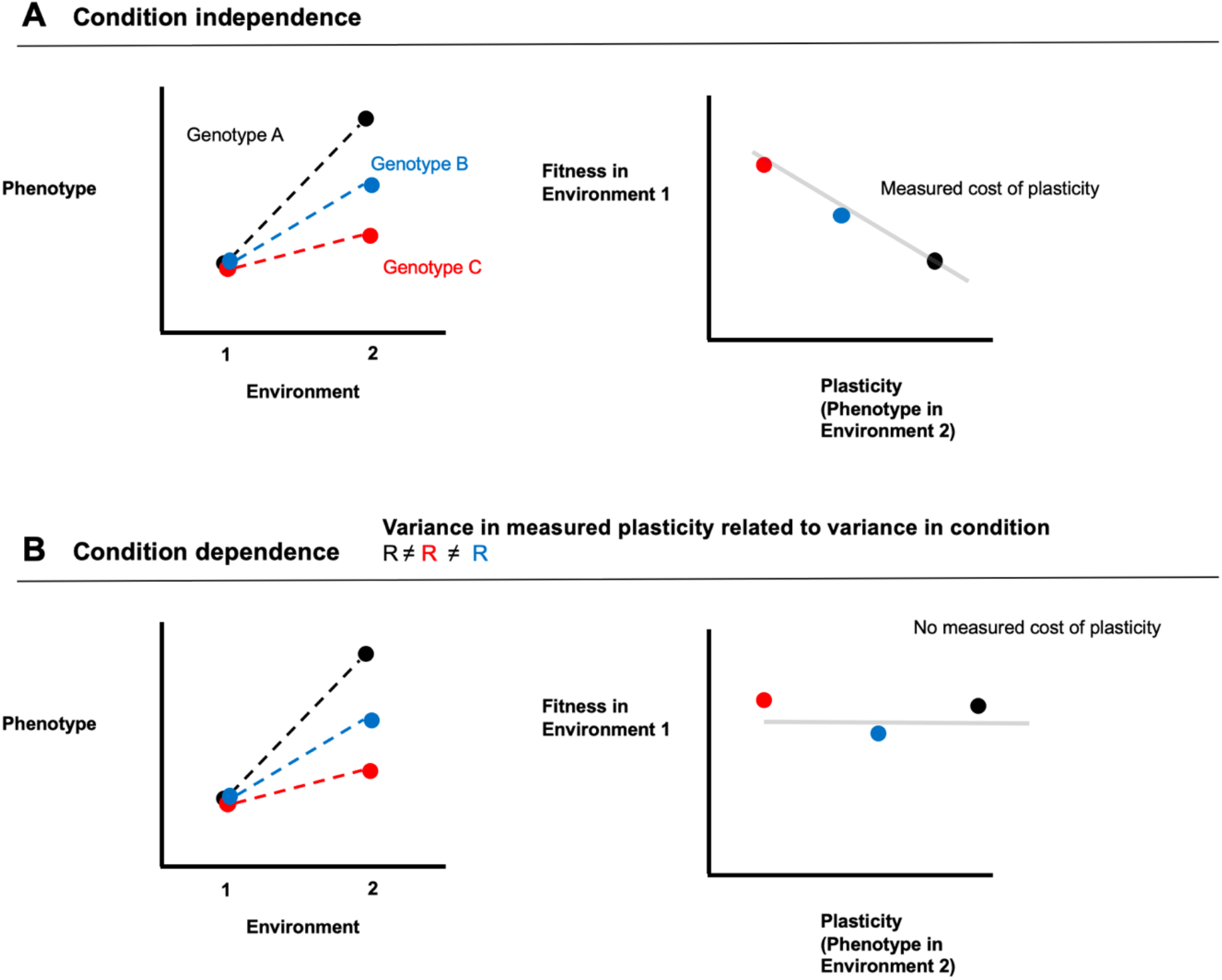
Panel A: illustrates the manifestation of a cost of plasticity when plasticity or trait expression is condition-independent, in which genotypes that have greater plasticity pay a fitness cost in the focal environment (controlling for trait expression in the focal environment) compared to less-plastic genotypes due to the cost of genotype plasticity. Panel B illustrates how fixed genotypic costs of plasticity can be masked by variance in condition. In this case, differences in trait expression in environment 2, and thus differences in phenotype plasticity between genotypes, are the result of variance in condition. In this case a cost of plastic resource allocation would not be measured at the level of trait expression, even if such a cost exists. In this figure we have illustrated the case where fitness effects of variation in trait expression in the focal environment have been controlled for; e.g., as would be the case in a multiple regression.

Thus, inference of costs of phenotypic plasticity depend upon the interpretation of covariance between fitness in one environment and trait expression in another. Yet such fitness costs of plasticity are rarely observed (Van Buskirk & Steiner 2009; Auld *et al*. 2010); often no fitness effects of phenotypic plasticity are found (Scheiner & Berrigan 1998; DiGiacopo & Hua 2020), or in some cases, the fitness effect is positive (Kenkel & Matz 2016) even after accounting for direct effects of trait expression in the focal environment. In other cases, costs are evident (Agrawal *et al*. 2002) or may depend on the degree of environmental quality (Van Buskirk & Steiner 2009).

This often “missing” cost of phenotypic plasticity represents an important unsolved paradox in evolutionary ecology (Snell-Rood & Ehlman 2021), and has led to a lack of clarity in the study of phenotypic plasticity and its role in adaptive evolution. On the one hand, it has been proposed that the empirical data suggest costs of plasticity may not exist or are very weak (Van Buskirk & Steiner 2009). On the other hand, some have suggested that costs may exist but simply be difficult to measure (Murren *et al*. 2015; Snell-Rood *et al*. 2018), and thus theoretical models and empirical studies of plasticity should still presume that costs of plasticity exist and play an important role in constraining reaction norm evolution. Indeed, theoretical models of the evolution of plasticity still rest on the assumption that there is a fitness cost that constraints the evolution of complete phenotypic plasticity (e.g., Chevin *et al*. 2010; Xue & Leibler 2018; Lande 2019; Ratikainen & Kokko 2019; Scheiner *et al*. 2020). In many ways, resolving the puzzle of missing costs of plasticity is essential to unifying theoretical and empirical studies of the evolution of plasticity, especially in light of new empirical approaches (Phillimore *et al*. 2010; Phillimore *et al*. 2012; Kenkel & Matz 2016; Rivera *et al*. 2021; De Lisle *et al*. 2022) that have reinvigorated the study of the evolution of plasticity.

The methods for detecting costs of plasticity may fail for a variety of reasons, all of which relate to unidentified sources of variation. For example, if unmeasured aspects of the genotype affect both plasticity and fitness in the focal environment, then costs may be underestimated or even overestimated. Likewise, if this shared effect varies between test conditions in the same population, then cost estimates will vary. Life history theory has addressed similar obstacles in detecting trade-offs among fitness components, beginning with the well-known work of van Noordwijk & de Jong (1986). Here we extend this approach to illustrate how variance in individual condition can mask costs of plasticity. We develop two models of condition dependence; in the first case, we explore the simple situation where the genetic component of plasticity carries a cost, leading to the evolution of condition dependent plasticity; this model is consistent with the expectation that costly traits under persistent selection may evolve to be condition-dependent. In the second model, we explore the case where costs of plasticity occur, but it is trait expression itself that is condition-dependent. We show both of these quite different forms of condition-dependence can readily mask fitness costs of plasticity. We also demonstrate how differences in the extent of assessed plasticity resulting from degrees of differences between focal and test environments can influence estimates of costs, particularly when trait expression is condition-dependent. This second factor may account for the observation that assessed costs are relatively common when the test environment is stressful (Van Buskirk & Steiner 2009; Snell-Rood & Ehlman 2021). Finally, we use these insights to suggest experimental designs that will maximize the detection of any existing costs of plasticity.

## Methods and Results

### Estimating Costs under Condition-independence

We begin by describing a causal model for fitness, and how a such a causal model of fitness is related to a statistical regression model based on trait data to infer costs of plasticity. We focus on the case of two environments. We assume individual fitness in a focal environment is a function of trait expression in that environment (natural selection) as well as the cost of having a plastic genotype,

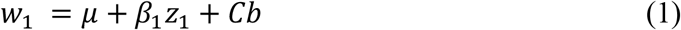

where *w*_1_ is individual fitness in environment 1 (the focal environment), *β*_1_ is natural selection in the focal environment, *z*_1_ is the trait value expressed in the focal environment, *b* is the fixed value for plasticity of the genotype, and *C* is the cost of such plasticity. We refer to *b* as “genotype plasticity” to be clear that it is a property of a genotype. Throughout, we refer to plasticity measured on actual traits as the “phenotype plasticity” to distinguish it from this true fixed cost. Equation 1 represents an assumed causal model for fitness effects of plasticity; noteworthy is that we have assumed fitness costs of plasticity are a fixed property of a genotype, depending only on its genotype plasticity *b* (expected to be the same for a given genotype across all environments) and the cost parameter *C* (a population parameter fixed across environments). Thus, we assume costs are fixed within a genotype and are paid across all environments, which is perhaps the simplest form of a cost of plasticity. We make no assumptions about selection on *z* in other environments. DeWitt et al. (1998) identified five non-exclusive mechanisms which may generate a cost of plasticity. These include maintenance costs (the cost of maintaining sensory and regulatory mechanisms), production costs, information acquisition costs, developmental instability, and genetic costs. Of these categories, maintenance, information acquisition, developmental instability, and genetic costs are all expected to often be a fixed property of the genotype, and so are consistent with the assumptions of Equation 1. We discuss environment-dependent production costs later.

Importantly, genotype plasticity *b* cannot be measured with trait data from only one environment, and so is instead typically inferred from trait data from the same genetic backgrounds expressed in two or more environments in a GxE design. In this approach, *b* is assumed to be proportional to phenotype plasticity; that is, trait expression in a another environment, *z*_2_. The cost of plasticity is thus assumed to be related to the covariance between fitness in one environment and trait expression in another, *cov*(*w*_1_, *z*_2_). Thus, with trait data from the same set of genotypes in two environments and fitness measured in one of the environments, the following regression has been proposed (Van Tienderen 1991; DeWitt 1998; Scheiner & Berrigan 1998) to infer costs of plasticity,

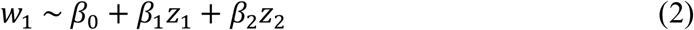

where the estimate 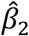 is interpreted as being an estimate of the cost of plasticity in equation 1, *C*. Note that a conceptually equivalent but more complex model could instead model *β*_2_(*z*_1_ − *z*_2_). Figure 1A shows an example of how a cost of plasticity is expected to manifest a reduction of fitness of a plastic genotype relative to a non-plastic genotype in a GxE experiment. Our goal is to understand when and why the regression model in equation 2 may fail to adequately describe the causal fitness effects assumed in equation 1, and to do so we need to develop more explicit descriptions of trait expression in each environment.

We first describe trait expression in two environments independent of condition,

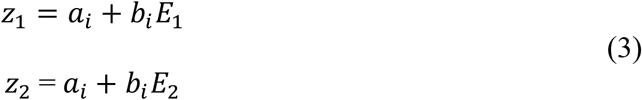

Where *b*_*i*_ is genotype plasticity or the reaction norm for genotype *i, E*_1_ the environmental value for environment 1 (for example, mean temperature), *a*_*i*_, the fixed (non-plastic) component of trait expression for genotype *i*.

When trait expression is plastic but condition-independent as described in equation 3, the cost parameter *C* is estimable in a multiple regression of the form

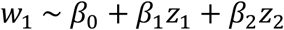

where the estimate 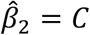 *if* all covarying traits affecting fitness are included in the multiple regression. Thus, in a GxE experiment where fitness data in at least one environment is combined with trait expression across two environments for a set of genotypes, fixed costs of plasticity for condition-independent traits can be inferred from such a multiple regression, as has been widely proposed and implemented (Van Tienderen 1991; DeWitt 1998; Scheiner & Berrigan 1998; Van Buskirk & Steiner 2009) and as illustrated in Figure 1A.

### Model I: Condition-dependent plasticity

Life history theory suggests that phenotypes that are costly to express will often evolve patterns of condition-dependent expression, where expression is higher in individuals that have the resources to pay the marginal costs of the phenotype (Rowe & Houle 1996; Houle & Kondrashov 2002; Lorch *et al*. 2003). Thus, if plasticity is costly yet also carries fitness benefits in variable environments, we may expect the evolution of condition-dependent plasticity in which plasticity is a function of individual condition,

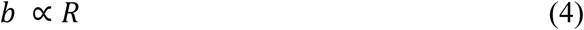

where *R* is condition; the total pool of resources an individual has available to allocate to phenotypes and fitness components (Figure 2A). In this model, we must also modify our fitness function to reflect the fact that condition *R* will itself affect fitness via other paths besides plasticity,

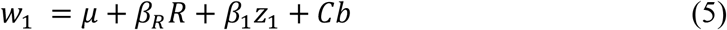

where *β*_*R*_ is the strength of selection on condition, independent of its effects on plasticity. This parameter reflects the summed effects of all the condition-dependent traits that affect fitness. In this fitness model, condition affects fitness both directly, and indirectly via effects on plasticity. We assume in Model I, for simplicity, that condition is a property of a genotype that is constant across environments. This general model of condition-dependent plasticity is illustrated in Figure 2A.

**Figure 2.**
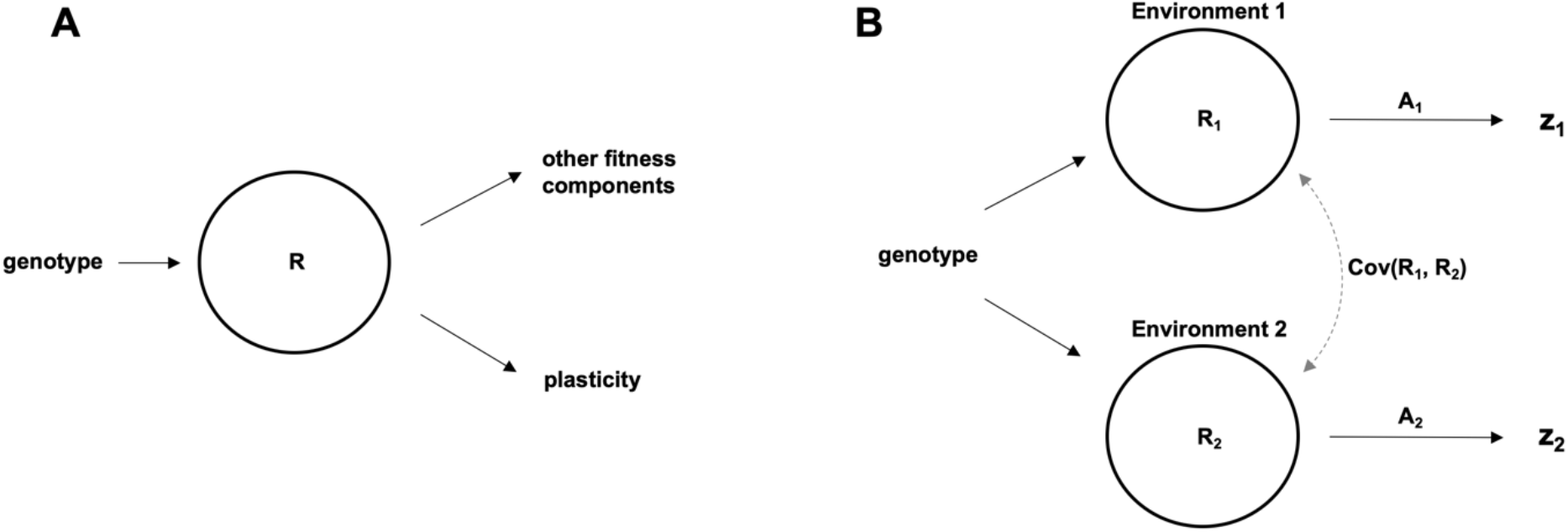
Two models for how condition may affect differential trait expression in multiple environments. In A, Model I, the total pool of resources available, *R*, which we call condition, is correlated with both fitness components and the degree of costly plasticity that is expressed. We note that this model is agnostic to the exact developmental causality of condition-dependence, and assumes only the existence of a relationship between these components. Variance in condition can mask costs of plasticity, because in this case individuals that have high plasticity (and thus pay a high cost) will nonetheless have high fitness despite the cost because condition also positively affects other fitness components. Panel B, Model II, represents a model of condition-dependent *trait* expression in two environments, where a condition-independent cost of plasticity may be found in any difference in resource allocation across in environments (A_1_ vs A_2_). Trait expression (z) is determined by condition (R), and allocation (A) in environment 1 and 2. Differences in resource acquisition and/or allocation across environments leads to differential trait expression across environments-phenotypic plasticity. In this model, we have assumed costs of plasticity arise when genotypes differ in resource allocation strategy (A) across environments. In this model, (co)variance in condition can mask costs of plasticity by generating variance in trait expression across the environments that is independent of variation in the costs paid. Note that for simplicity, we have not expanded these path diagrams to directly compare the complete path to fitness; rather to illustrate the differing roles of condition.

We can understand the effects that variance in condition has on our inferences of costs by expanding *cov*(*w*_1_, *z*_2_), the relationship between fitness in one environment and trait expression in another, that has typically served as the basis for inferring costs of plasticity. Assuming no covariances with *a* (the fixed genetic component of trait expression) for simplicity,

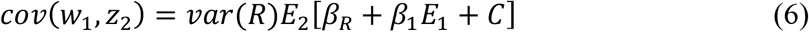

Equation 6 shows that fitness effects of plasticity, *cov*(*w*_1_, *z*_2_), are fundamentally influenced by variance in condition (see Figure 1B). This covariance will be negative, thus implying a net cost of plasticity in the focal environment, only when

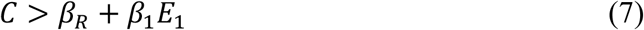

where *C* is in absolute terms. Although the second term on the right hand side of equation 7 can be controlled for in a multiple regression that includes *z*_1_, even in this case the cost of plasticity must be greater than total selection on condition itself (*β*_*R*_) in order for a negative cost to be inferred, an unlikely situation. When *β*_*R*_ is greater than the cost of plasticity, a positive fitness effect of plasticity will be inferred despite the existence of a cost, even when controlling for *z*_1_, and the magnitude of this positive fitness effect will be proportional to the variance in condition *var*(*R*), as illustrated in Figure 1B. In the appendix, we show that this model of costly plasticity can alternatively be framed as a specific case of the classic tradeoff model of van Noordwijk & de Jong (1986).

This simple model shows that when plasticity is itself condition dependent, inferring a cost of plasticity in a GxE experiment will be difficult or impossible unless variation in condition or resource acquisition can be controlled. This model of condition-dependence can thus readily explain the variable and weak costs of plasticity that have typically been inferred in previous experiments. However, it cannot immediately explain the finding that costs are typically inferred to be greater in stressful or poor quality environments (except to the degree that such manipulations affect within-environment variance in condition), and represents only one way that condition may impact variance in trait expression across multiple environments.

We have introduced this simple model, a special case of van Noordwijk & de Jong’s general life history model, to illustrate a likely explanation for why costs of plasticity are rarely observed. However, it is noteworthy that this model deviates substantially from the casual fitness model envisioned by past workers measuring costs of plasticity (DeWitt 1998), in that we have assumed that condition affects fitness independently of both trait expression and plasticity. Although life history theory and empirical data support such a general fitness model (van Noordwijk & de Jong 1986; Rowe & Houle 1996), it is also true that any discrepancy between the assumed fitness model and the actual causes of fitness (due to missing traits, microenvironmental covariances, etc.) can result in misleading estimates of a cost to plasticity (Morrissey *et al*. 2010). Next, we present an alternative model of condition dependence that shares the same causal fitness model typically assumed in studies of plasticity cost, but includes a condition effect on the measured trait, but no independent effect of condition on fitness.

### Model II: Condition-dependent traits

In our second model we assume that fitness is caused solely by the trait and by plasticity, as in equation 1,

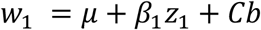

with no independent causal path for condition. Thus, this causal model of fitness is the same as that assumed in typical analyses of the cost of plasticity. We now assume trait expression is the result of both plastic resource allocation and the total pool of resources an individual has available to allocate, or condition. Thus we have assumed the trait is costly, in terms of resources, to express. We can modify our description of trait expression accordingly,

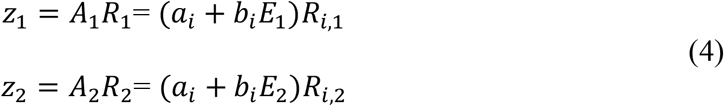

where *A*_1_ is the pattern of resource allocation in environment 1 containing both plastic and fixed components, and *R*_*i*,1_ is condition of individual *i* in environment 1, with a corresponding term for environment 2. In this model, condition affects fitness only indirectly via its effects on expression of *z*. Variation in condition may be genetic or non-genetic. Importantly, in this model two separate components contribute to the phenotypic plasticity (that is, *z*_2_): 1) plastic changes in resource allocation *A* across the environments determined by *b* (which we have assumed is costly, as stated in equation 1), which can be described as the ability to match allocation strategy to the environment, and 2) plasticity arising from variation in resource acquisition *R*, or condition (which we have assumed is cost-free), which is simply the amount of resources an individual has available to allocate to traits. As we show below, if a substantial amount of variation in plasticity is determined by this second component, variation in condition, then costs will be difficult to infer even if they exist.

This general model of trait expression is shown in Figure 2B, and can be seen as an extension of previous hierarchical acquisition/allocation models (van Noordwijk & de Jong 1986; Houle 1991; Rowe & Houle 1996), developed in the context of life history evolution, to the case of resource acquisition in two distinct environments. Our model is thus conceptually similar, although very different in purpose, to that developed by Zajitschek and Connallon (2017) in the context of separate sexes instead of separate environments. Figure 2 illustrates some of the conceptual difference between our Model I and Model II. In Model I, we assume that all variation in plasticity is costly, and variation in plasticity is condition-dependent. In Model II, we assume condition-dependent trait expression in that trait expression is a function of condition and allocation, where phenotype plasticity arising from differences in condition across environments carries no cost, while plasticity arising from differential resource allocation across environments does carry a cost. In Model I, condition was assumed to have independent effects on fitness; in Model II condition only affects fitness via z.

As before, we can expand *cov*(*w*_1_, *z*_2_) to understand how variance in condition affects inference of costs in model II. Under this case of condition dependent trait expression (following Bohrnstedt & Goldberger 1969; see Appendix), using equations 1 and 4 this covariance will be negative (and thus suggest a net cost of plasticity) when

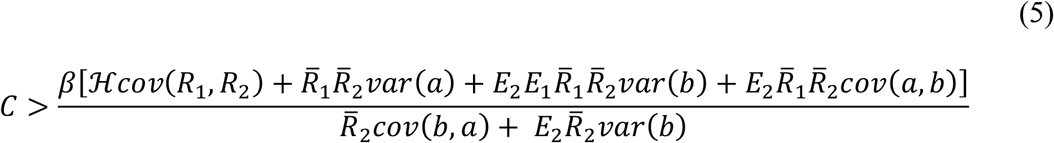

For simplicity assuming no covariance between *a* or *b* and *R* and where 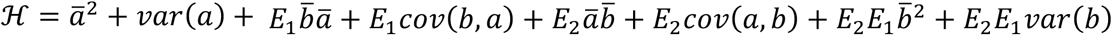. Although this expression is complex, it illustrates that fixed genotypic costs of plasticity have to be very high to manifest a negative *cov*(*w*_1_, *z*_2_), particularly when *cov*(*R*_1_, *R*_2_) is positive. We can see this in Figure 3A, noting the very high magnitude of C needed to generate negative *cov*(*w*_1_, *z*_2_). Because this magnitude can be interpreted relative to the strength of selection, this suggests that biologically-realistic values of a cost to plasticity will not generate negative relationships between trait expression in one environment and fitness in another, unless covariance in resource acquisition is almost perfectly negative across the two environments or there is no phenotypic selection (i.e. all fitness variance arising from costs of *b*).

**Figure 3.**
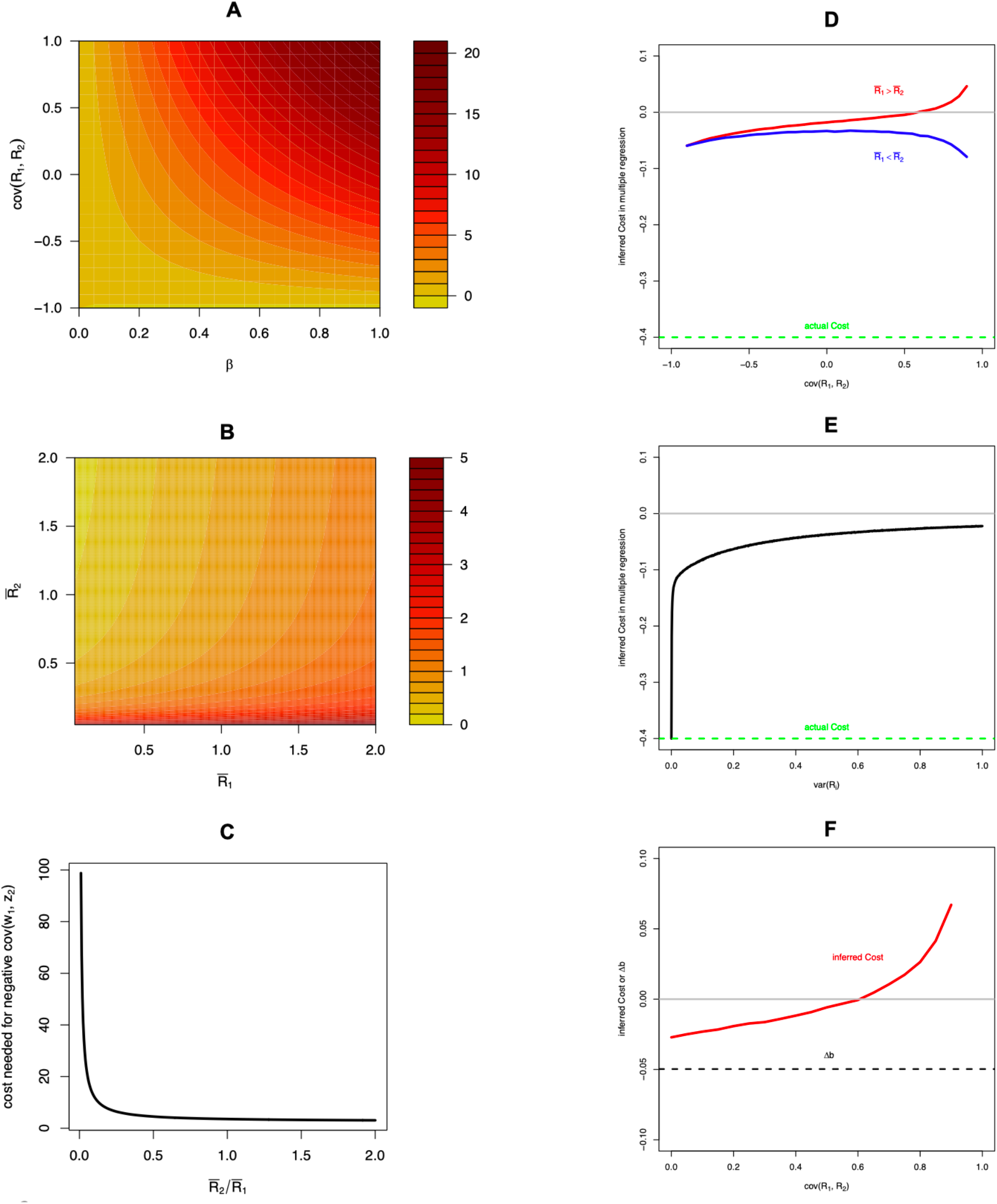
Effects of variation in condition on inference of plasticity costs under Model II, condition-dependent trait expression. Panels A-C show minimum values of plasticity cost that are required to generate a negative covariance between trait expression in one environment and fitness in another, *cov*(*w*_1_, *z*_2_). Panels D-F show inferences cost costs in a multiple regression controlling for the direct fitness effect of traits. Panel A: Strong positive genetic covariance in resource acquisition across environments makes it more difficult to detect costs, as the cost of plasticity must be higher to result in negative *cov*(*w*_1_, *z*_2_). Parameter values, 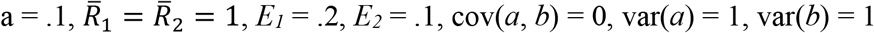. Panels B and C illustrate the effects of average condition in each environment. Parameter values,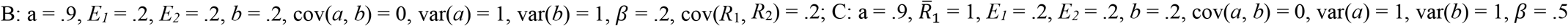, cov(*R*_1_, *R*_2_) = .2. In all panels D-F, inferred cost is the partial regression coefficient 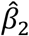 from the multiple regression model *w* ∼ *β*_0_ + *β*_1_*z*_1_ + *β*_2_*z*_2_. Panel D shows the inferred cost as a function of the covariance in resource acquisition, for the case where the focal environment is poor relative to the second environment (blue) and the case where the focal environment is high quality relative to the second environment (red). Panel E shows inferred cost as a function of the variance in resource acquisition (assumed equal across environments, with zero covariance). Panel F shows the case where costs of plastic resource allocation are high (-0.6), natural selection on the trait is weak, and covariance in resource acquisition is high. Dashed black line in panel F shows the expected evolutionary change in the allocation reaction norm in the focal environment, *cov*(*w*_1_, *b*). Green dashed line in panels D and E show the actual cost of the allocation reaction norm *b*, which was omitted in panel F for scale. In Panel D inferred costs were calculated as the average of 1000 partial regression coefficients computed numerically from a random sample of 100 individuals with 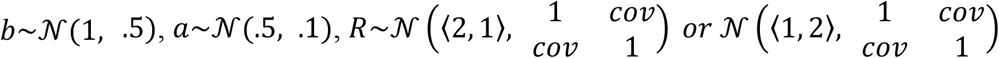, for each value of *cov*, assuming *β* = .5, *E*_1_ = 1, and *E*_2_ = 2. Calculations in panel E were equivalent except 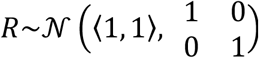. In Panel F, *β* = .2.

Importantly, Model II shows that when trait expression is condition dependent across two environments, the observed degree of phenotypic plasticity in the trait itself will be in-part condition dependent, and variance in phenotypic plasticity will be in part determined by (co)variance in condition across the two environments. If a measure of individual condition, *R*, is available, or a measure of *b* controlling for condition, then an appropriate multiple regression could be fit to obtain an estimate the cost parameter *C*. However, such measures are rarely available and not typically used as control variables in previous GxE studies aimed at inferring costs of plasticity. For example in a multiple regression of the form

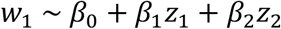

(DeWitt 1998) any variance in condition *R* will affect both the measure of plasticity (*z*_2_) and fitness in the focal environment (*w*_1_), resulting in a biased estimate of the true cost of plasticity such that 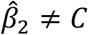. This effect of variance in condition on cost inference in a multiple regression is illustrated in Figure 3D-F; (co)variance in condition masks inference of costs in a multiple regression, regardless of the nature of environmental quality (Figure 3D, E). Whenever variance in condition exists, this variance generates variance in the phenotype plasticity that is independent from costs associated with genotype plasticity, and so this variance masks costs (Figure 3E; see also Figure 1B). In the extreme, when covariance in condition across environments is strong, costs of genotype plasticity are high, and the strength of natural selection on the trait in the focal environment is weak, it is possible for (co)variance in condition to lead to the inference of positive fitness effects of plasticity even when the expected evolutionary response of genotype plasticity is negative (Figure 3F). This extreme scenario suggests it is possible to infer, from a multiple regression analysis of trait data in a GxE design, direct selection for increased phenotypic plasticity when the evolutionary response of genotype plasticity is in fact negative. Although it is unclear how frequently such conditions occur, in general these effects (Figure 3D-F) illustrate that costs of plasticity incurred within the path from genotype to trait expression will be readily masked by condition, a point also made in simpler visual terms in Figure 1. Although we have assumed a cost of at the level of resource allocation, assuming instead that a cost is incurred through differential resource acquisition would lead simply lead to a reversal of the expected effects of acquisition and allocation on *cov*(*w*_1_, *z*_2_).

Equation 5 also illustrates that average resource acquisition in each environment influences ability of costs to be inferred. In particular, costs of plasticity are most readily inferred when the quality of the focal environment 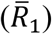 is low relative to environment 2 (that is, when 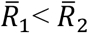), and the interaction between average resource acquisition in the two environments determines whether allocation costs generate *cov*(*w*_1_, *z*_2_) (Figure 3). These effects can be illustrated more simply in Figure 4, which shows how changes in mean condition in environment 1 affect fitness, while changes in mean condition in environment 2 affect the inference of plasticity, and so average condition across the two environments interact to influence *cov*(*w*_1_, *z*_2_).

**Figure 4.**
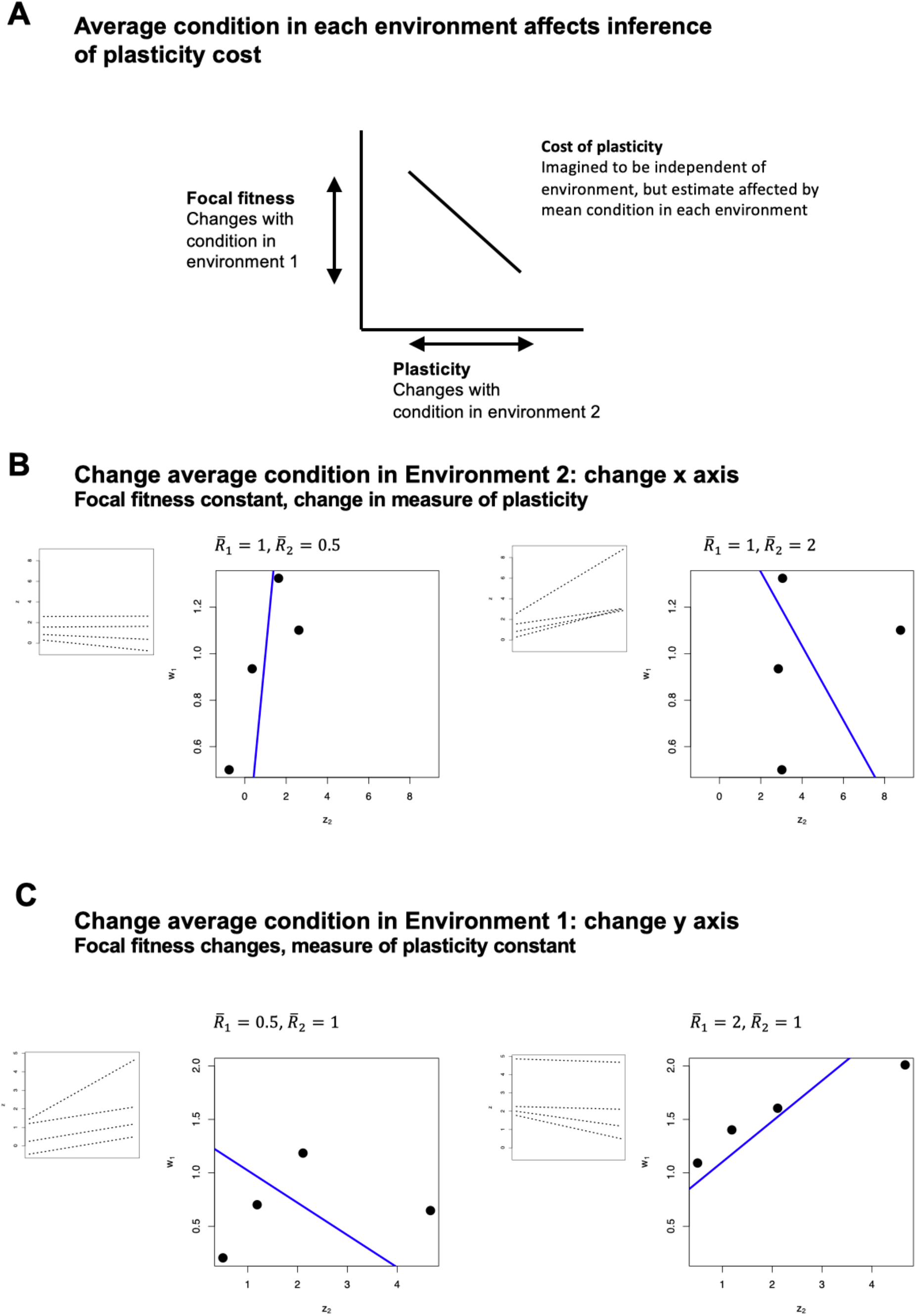
Quality of both environments determines inference of costs of plasticity under Model II, where trait expression is condition-dependent. Panel A shows how average condition in the focal and second environment affect fitness in the focal environment and the estimate of plasticity, respectively, which are the y and x axes determining *cov*(*w*_1_, *z*_2_), which is used to infer the cost of plasticity. Panel B shows how changes in mean condition in Environment 2 affect this covariance; under a constant fixed cost of plasticity and assuming the same sample of genetic variants for b, a, and 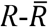, increasing 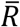 in Environment 2 leads to the inference of more costly plasticity. Panel C shows the affects of changing 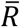 in the focal environment for the same set of genetic variants as in Panel B; increasing 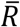 in Environment 1 makes costs more difficult to reveal. Inset panels show individual norms of reaction for each genotype. Blue line is the partial regression coefficient for *z*_2_ from a multiple regression controlling for *z*_1_.

### Environment-dependent costs

We have explored the case where costs of plasticity are a fixed property of a genotype, paid in all environmental contexts, although inference of these costs may depend on the environmental conditions (as shown above). Alternatively, costs of plasticity may only be incurred when a plastic trait is actually expressed. For example, a “production cost” (DeWitt *et al*. 1998) of a plastic phenotype, where the costs relative to a fixed strategy are only paid when the trait is produced. Statistically, we could define such a cost as an interaction for fitness between the plasticity of a genotype and the environment,

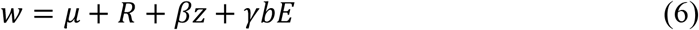

Where *γ* defines the fitness cost of plasticity *b* that only manifests in certain environmental contexts *E* (the same environment in which fitness is measured). This cost, *γ*, could in principle be inferred if one had estimates for *w*, R, z, b across multiple environments for a set of genotypes. However, the challenges produced by condition dependence, shown above for the much simpler case of fixed costs, will only be greater in this more complex case. As an example, consider the case of Figure 4B, C. In these cases, where costs are in fact fixed, an analysis of phenotype data across the different environments would suggest costs to differ across these environmental contexts. Note also that Model I could be modified so that *R*_1_ ≠ *R*_2_, which would lead to similar complexities of costs that vary across environments.

## Discussion

Phenotypic plasticity is expected to be costly, as without costs or some other constraint organisms would often evolve to be completely plastic, which they do not. Costs of plasticity are also expected on the simple basis of investment into mechanisms required to shift developmental trajectories or behavior in response to the environment, or as a result of imperfect environmental cues (DeWitt *et al*. 1998; Snell-Rood *et al*. 2018; Snell-Rood & Ehlman 2021). These intuitive arguments for the prevalence of a cost of plasticity are at odds with the empirical evidence for such costs. Quantitative genetic GxE experiments in a variety of plants and animals (Van Buskirk & Steiner 2009; Auld *et al*. 2010), as well as more recent analyses of gene expression data (Kenkel & Matz 2016), have revealed fitness effects of plasticity to be weak, nonexistent, or sometimes even positive. This apparent puzzle of missing costs of plasticity has been left largely unsolved (Murren *et al*. 2015; Westneat *et al*. 2019; Snell-Rood & Ehlman 2021), all while study of the evolution of plasticity has advanced via new empirical and theoretical approaches that are ultimately based on the presumption of existence of costs. Here, we show that condition-dependence may generally mask costs of plasticity. Using two models of condition dependence in multiple environments, we show that the paradox of missing plasticity costs can be readily explained when plasticity or trait expression is dependent on condition. In both cases, phenotypic plasticity itself is condition dependent, and fitness costs incurred by plastic genotypes can be masked by variance in individual or mean condition. Our analyses provides an explanation for why costs of plastic trait expression are rarely inferred in GxE experiments, even if fitness costs of plasticity do exist. Our work also illustrates that the interpretation of selection coefficients in GxE experiments may be far less straightforward than has typically been appreciated, even for the simplistic cases we have explored where costs of plasticity are a fixed property of a genotype paid in all environments.

Our models can account for several features of empirical evidence for costs of plasticity. First, our finding that variance in plasticity generated by variance in condition masks costs of plasticity is consistent with the empirical finding of low or nonexistent costs of phenotype plasticity (Van Buskirk & Steiner 2009; Auld *et al*. 2010). Indeed, both of our models show that this masking effect can even lead to the inference of positive fitness effects of phenotype plasticity even when costs of plasticity exist. Second, our model of condition-dependent trait expression (Model II) may account for the finding of elevated costs of plasticity inferred in stressful environments (Van Buskirk & Steiner 2009). Our analysis shows that average condition in both the focal and second environment interact to determine whether a cost of plasticity can be revealed from trait data. When average condition is low (that is, environmental quality is low) in the focal environment relative to the second, costs are more readily revealed. This is because such a combination results in low fitness in the focal environment in combination with a high degree of inferred plasticity; in contrast, high average condition in the focal environment relative to second environment generates a scenario with high mean fitness and a low overall measures of plasticity, thus making costs more difficult to reveal even if they exist (Figure 4). Finally, our models may account for the finding that costs of plasticity are more readily revealed in analysis of recombinant inbred lines than in analysis of naturally-sampled genetic families (Callahan *et al*. 2005; Dechaine *et al*. 2007; van Kleunen & Fischer 2007). Theoretical models of the evolution of condition dependence indicate that one way in which covariance between condition and trait expression can develop as a result of disequilibria between alleles influencing the two (Lorch *et al*. 2003) or through the spread of unlinked modifier alleles (Houle & Kondrashov 2002), and so generation of recombinant inbreds would be expected to reduce this covariance, making costs of plasticity more easily revealed.

Together, these results suggest that care should be taken in experimental design to restrict within-environment variance in condition, or resource acquisition (to the extent this is possible) and to contrast environments that differ in quality, if the aim of the experiment is to infer costs of plasticity. Alternatively, GxE designs can be combined with other approaches, such as experimental evolution, to understand the nature of costs of plasticity. For example, Brennan et al. (2022) combined experimental evolution and a reciprocal transplant to reveal the likely existence of costs of transcriptional plasticity in marine copepods.

Empirical evidence suggests that both forms of condition-dependence that we have explored may be commonplace. In general, many of the traits that have been the focus of previous studies of plasticity are traits that are closely related to life history and components of fitness. This includes traits such as the timing of reproduction or maturation, growth rate or size at maturity, body size, and defense against natural enemies (Agrawal 2001; Relyea 2002; Phillimore *et al*. 2010; Charmantier & Gienapp 2014); traits that are closely related to life history and thus are expected to evolve condition dependence (Rowe & Houle 1996). Plasticity itself has also been shown to exhibit condition dependence, particularly in the case of mating and display traits (De Lisle & Rowe 2014; Kuczynksi *et al*. 2016; Ziegler *et al*. 2018; Anichini *et al*. 2019).

As a caveat, our assumed models for fitness are simplistic, particularly in model II where we assume directional selection in the focal environment (DeWitt 1998), and make no explicit assumptions about selection in other environments. Our numerical exploration of Model II (Figures 3, 4) explored the specific case of positive directional selection in the focal environment; consistent with the assumption of a costly trait, condition dependence is expected to evolve in such circumstances (Rowe & Houle 1996; Houle & Kondrashov 2002; Lorch *et al*. 2003). Moreover, in this model we assumed that condition can only affect fitness via effects on expression of a single trait. A more realistic and general causal model of fitness would perhaps include stabilizing selection towards an optimum, and multiple pathways in which condition can affect fitness independent of *z*. However, our development of simplistic fitness models was purposeful; our aim was to understand how condition dependence can influence inference of plasticity costs even when the true causes of fitness are similar to (Model I) or identical to (Model II) that envisioned in past studies (e.g., DeWitt 1998).

Others have suggested that resource acquisition and condition dependence, as well as missing traits that affect fitness, may all make it difficult to measure costs of plasticity (Buchanan *et al*. 2013; Murren *et al*. 2015; Snell-Rood *et al*. 2018). Our analysis demonstrates explicitly why this is so and the degree of the problem. When trait expression and plasticity are condition-independent, costs of plasticity can indeed be measured in a phenotypic multiple regression. When plasticity is itself condition-dependent, or when trait expression is condition dependent and costs of plasticity arise through changes in resource allocation independent of condition, the effects of such costs on covariance between phenotype and fitness are masked by phenotypic variance that arises from differences in condition. In this case, the problem is more than one of missing traits; without some separate measure of condition, inferring costs of plasticity will always be biased by variance in resource acquisition. Our results are also consistent with the model of Matthey-Doret et al. (2020), who show using a model of gene networks that alternative developmental mechanisms can allow cost-free plasticity to emerge.

We have shown that costs of plasticity may be especially hard to infer when the cost is paid at some point along the path from genotype to phenotype, rather than at the final level of phenotype expression itself. We believe that most previous ideas surrounding the causes of potential costs of plasticity fit this assumption, that costs are paid on one of several components of the path from genotype to phenotype. For example, DeWitt et al. (1998), who lay out 5 non-exclusive categories of costs of plasticity, propose that these costs likely arise somewhere on the path from detection of environment → information processing → regulatory mechanism → production machinery → trait expression. Our model shows that even when costs are high at any one point in the path, variance at another point that is cost-free will mask the cost when measured at the level of trait expression.

If plasticity is adaptive, an open question is how genetic variance in plasticity is maintained (which appears to often be the case; Scheiner 1993) in the face of persistent natural selection. Our model shows that if trait expression in multiple environments is condition-dependent, then expression of plasticity will itself be condition dependent and a portion of the standing variance in a phenotypic plasticity will reflect standing (co)variance in condition across environments. Because we define condition as the total pool of resources an individual has available to allocate, we expect condition to be a large mutational target, and thus capture novel mutational input across the genome (Rowe & Houle 1996). Thus, our model suggests genic capture may provide a mechanism maintaining variance in adaptive plasticity.

We have attempted to provide some closure for the unsolved problem of costs of plasticity. In many ways, the study of phenotypic plasticity has moved on from the uncertainty surrounding costs that reached a zenith over a decade ago (Auld *et al*. 2010). Yet the issue remains, and although the field has moved on two segregating viewpoints linger: do costs exist, but are hard to measure? Or are costs simply so weak as to be unimportant? We show, by recasting the problem of plasticity as one of differential resource acquisition and allocation, why costs may indeed be prevalent and important but difficult to reveal.

## Acknowledgements

We thank Erik Svensson and his lab group for discussion and feedback. Funding was provided by the Swedish Research Council (grant 2019-03706), Formas (grant 2021-01096), and the Royal Physiographical Society of Lund (grants 42305, 41593) to S.P.D.

## Appendix

### Model I as a specific case of van Noordwijk & de Jong

We can rewrite Model 1, Figure 2A as a tradeoff between plasticity and component fitness,

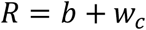

Where *w*_*C*_ are all fitness components affected by condition. We can further define the fraction of resources allocated to plasticity as *B*, where *b*_*i*_ *= B*_*i*_*R*_*i*_ and *w*_*C*_ *=* (1 − *B*_*i*_)*R*_*i*_. Following van Noordwijk & de Jong (1986; see their equation 5), we can define the covariance between fitness components and plasticity,

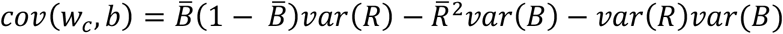

and note that this covariance will be positive (suggesting no cost) when the variance in condition is high relative to the variance in allocation strategy *B*. That is, when most of the variance in plasticity is due to variance in condition, as opposed to variance in how condition is partitioned into trait expression, costs of plasticity will be masked. The relationship between this version of Model I and the version presented in text can be seen by noting that in text we solve for *cov*(*w, z*_2_) under the assumption that *var*(*B*) *=* 0.

### Detailed derivation of equation 5

We can expand *cov*(*w*_1_, *z*_2_) under the assumptions of our Model II,

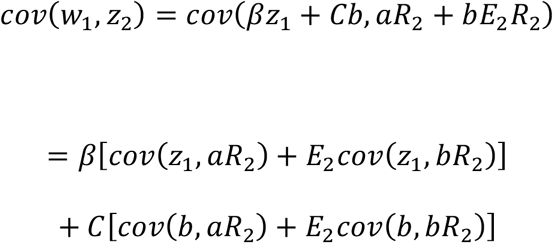

Which we can further expand assuming multivariate normality (for simplicity) following Bohrnstedt and Goldberger (1969),

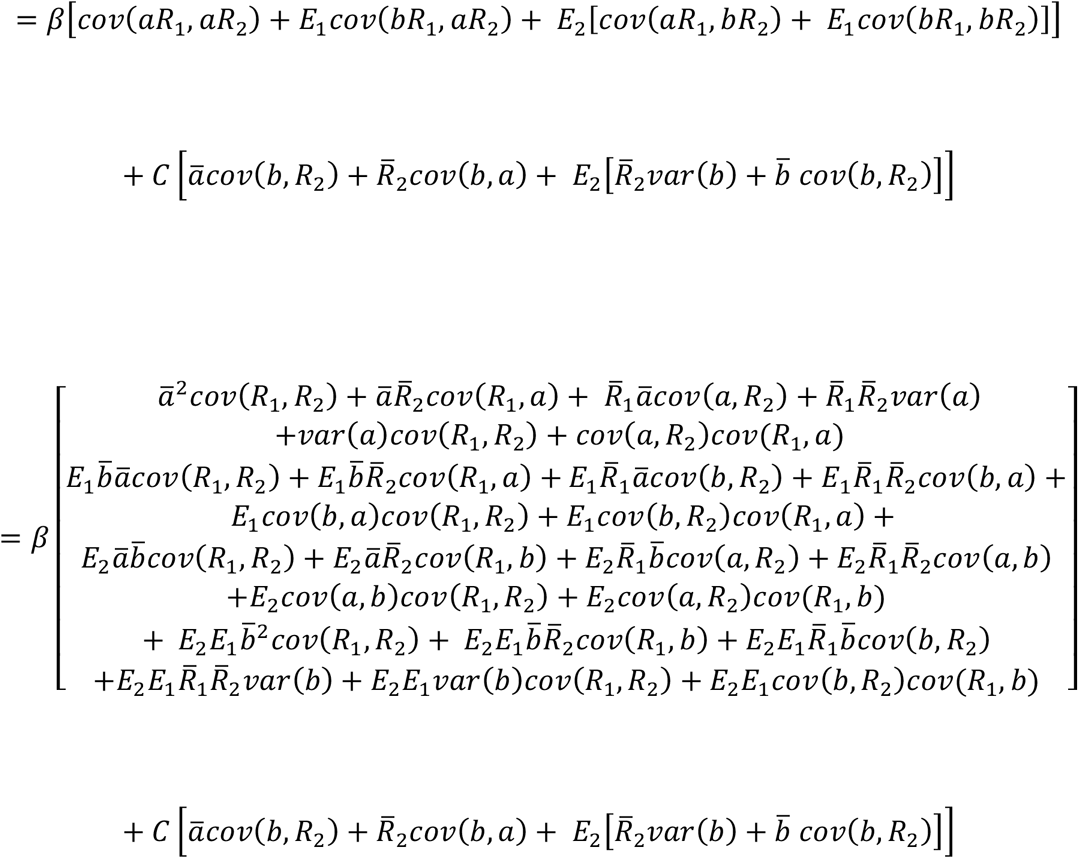

